# Extending and repeating assessments is a key tool to reveal barriers to effective management of protected areas

**DOI:** 10.1101/2020.11.26.399618

**Authors:** Marcos Eugênio Maes, Eduardo L. Hettwer Giehl, Natalia Hanazaki

## Abstract

Here we assessed protected area (PA) management effectiveness while also checking for potential barriers. We addressed 21 PAs of the State of Santa Catarina, southern Brazil. Out of these, we assessed 10 PAs under state level administration that lacked thus far a standardized assessment. For PAs under federal administration, we got assessment data from the government itself. We contrasted the PAs regarding the level of administration and collected a list of aspects that could result in differences in management effectiveness between PAs. We checked the relationship between PA aspects and mean effectiveness with linear models. The same aspects were also related to PA management elements, namely context, planning, inputs, processes, and outputs, using redundancy analysis. Management effectiveness and scores of management elements were found to be lower for PAs either with unresolved land tenure, lacking management plans, being under many pressures and threats, or being under state level administration. Our results call for extended assessments to reach the reality of the Brazilian PA network with different administration levels. Assessments must be carried out regularly since it is the only way to effectively flag a barrier, clear it, then find the next one to be tackled.

## Introduction

Creating protected areas (PAs) is the single most important strategy in conservation biology, even though many barriers limit the fulfillment of PA conservation aims. PAs are key for biodiversity conservation [1] by both reducing threats to biodiversity [2] and maintaining ecosystem services [3]. Global efforts toward biodiversity conservation have raised the coverage of PAs to ~15% in land areas [4]. Still, habitat loss and biodiversity decline are ongoing inside most PAs [5]. Such losses indicate PAs have been unable to meet all the goals they had been created for, a fact, at least in part, related to management ineffectiveness [6].

To aid in reaching successful management, the International Union for Conservation of Nature (IUCN) directed efforts for the systematization and assessment of PA management effectiveness. Such efforts produced a framework based on the principle of adaptive management [7] and the assessment of six elements: context, planning, inputs, processes, outputs, and outcomes [8]. Following this framework, different methods have been elaborated for assessing PA management effectiveness. Of these methods, the Rapid Assessment and Priority of Protected Area Management is often used (RAPPAM; see [9]). Thus far, management assessments cover only a small fraction of the global PA network [6] and indicate just a moderate effectiveness of ~50%, which tends to be even lower in developing countries [10]. In Brazil, assessments of management effectiveness have been carried out for PAs administered by the federal government; however, a substantial part of the Brazilian PA network is run under state level administration.

Understanding which PA’s aspects affect management results and directing actions toward the most relevant outcomes underlies effectiveness assessments and adaptive management, thus directing scarce resources toward conservation gains. In addition, PA management effectiveness depends on aspects like the size and qualification of staff and available funds. Such aspects are often dependent on administration levels--in the federal or state spheres--and public appeal, and here we checked their influence in PA effectiveness. Specifically, we postulate that there is an improvement in effectiveness for PAs under federal administration [11] and more stringent protection categories [11]; with established management councils [12] or including co-management by relevant partners [13]; with large staffs and funds [14]; with management plan [15] and with substantial public appeal indicated by attracting many visitors [16]. We also expected well-conserved ecosystems [17] to show more effective management results, because of both fewer pressures and threats to biodiversity [18]. Finally, land tenure resolution can also lessen many pressures and threats and thus lead to successful management [19].

Here we estimated the relative importance of aspects that can lead to distinct management effectiveness and, thus, for PAs to achieve their goals of biodiversity conservation. While the main aim of the study was in part exploratory, we tested the direction of the effects for each aspect as indicated above and expected financial and human resources to be the main drivers of management effectiveness of PAs. To test these hypotheses, we considered protected areas located in the State of Santa Catarina, under federal level and state level administration.

## Material and methods

### Protected area network

We assessed a PA network located in the Atlantic Forest domain, a region placed fourth for worldwide conservation priority [20]. Despite the threats that contribute to placing the domain under such status, the State of Santa Catarina (southern Brazil) has a high conservation potential with ~30% of native vegetation cover remaining, even though vegetation cover varies across the State [21]. The State has 99 PAs in several protection categories are under federal level (16 PAs), state level (10), municipality level (18), and private (56) administration (updated from [22]). Because of such differences across the PA network, management effectiveness is expected to differ, providing grounds for checking which aspects lead to better outputs.

### Assessment of management effectiveness

For the assessment of effectiveness, we selected 11 federal and 10 state level PAs out of the entire PA network of Santa Catarina. We assessed management effectiveness using RAPPAM [9] because it allows a standardized comparison between different contexts [10] and emphasizes effectiveness alongside pressures and threats. Pressures are actions that have negatively affected PAs in the five years before the assessment. In turn, threats that are persisting negative actions in the past that are likely to go on over the following five years. Effectiveness assesses whether the management is leading the PA towards its aims and considers the elements of context, planning, input, process, and outputs (outcomes are less related to management and were not assessed). RAPPAM uses several questions to characterize pressures and effectiveness. Combining scores assigned to effectiveness questions results in a management effectiveness index, which can be classified as low (<0.4), average (≥0.4 and <0.6) or high (≥0.6).

For federal level PAs, we used the most up to date RAPPAM assessment that was collected in 2015 [23] and covered 11 of the 16 federal PAs in the State of Santa Catarina. For state level PAs, which never underwent such assessment, we applied the whole RAPPAM to PA managers and, in addition, only the effectiveness questionnaire to a person of the State Environmental Agency and to two representatives of the management council of each PA. For state level PAs with either no council or when councilors could not be reached, we interviewed representatives of entities that were likely to take part in the council once it was established. Specifically, criteria for selecting representatives were a stated consent to take part in our study, basic knowledge of ecology or related fields, and enough knowledge about the PA. For each State PA, we calculated the effectiveness index as the median of the four questionnaires. This study was authorized by the Ethics Committee of Universidade Federal de Santa Catarina (53996516.7.0000.0121).

### Relating aspects of protected area with management outcome

We related potential drives with effectiveness in two ways to explain changes on either the effectiveness index or effectiveness elements. In both cases, we built a list of 13 PA aspects that could be responsible for changes to management outcome (e.g. PA administration level, protection category, funds and staff size, and so on; full list in Table 1). First, we built general linear models with aspects as predictor variables and PA effectiveness index as the univariate response. To reduce multicollinearity, we fit preliminary models and removed predictors with variance inflation factors (VIF) ≥ 4. Next, we choose which aspects were most strongly related to management effectiveness using a step-by-step removal process. This process was guided by values of corrected Akaike information criterion for small samples (AICc) and we stopped selection when we reached the lowest AICc-values. We used a multivariate approach to inspect for changes to the six elements underlying the effectiveness index. In this approach, the same 13 PA aspects were used as predictor variables in a Redundancy Analysis (RDA) with elements of effectiveness as the multivariate response. To check for multicollinearity, we fit a preliminary RDA and removed predictors with VIF ≥ 10. Then we reduced the full model in a step-by-step process using the highest adjusted R^2^ as the stopping criterion. We checked the overall predictor-response relationship of RDA and of each of the remaining predictors using permutation tests (9999 iterations). Finally, we also carried these analyses for the subset of State level PAs.

**Table 1.**
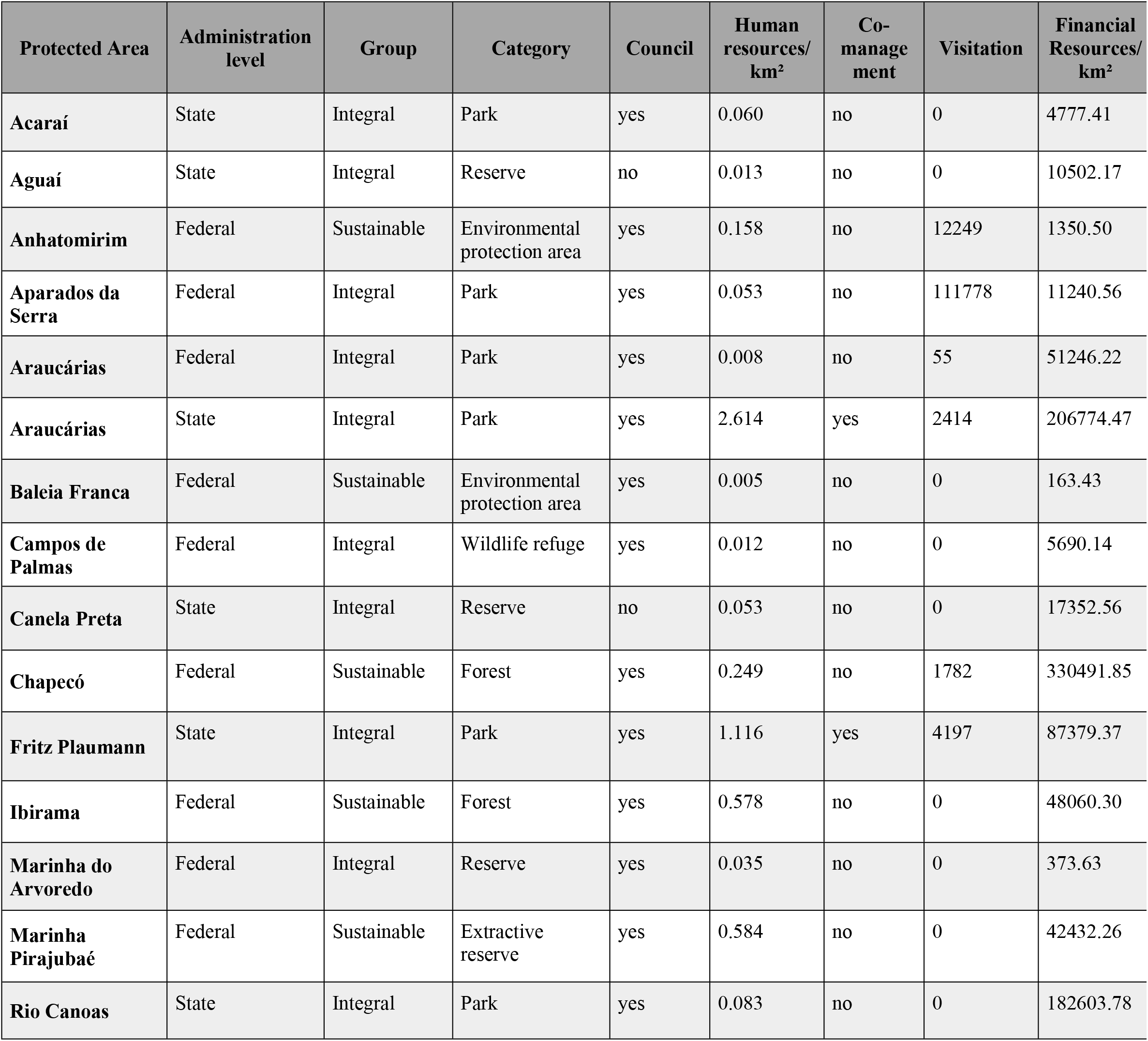
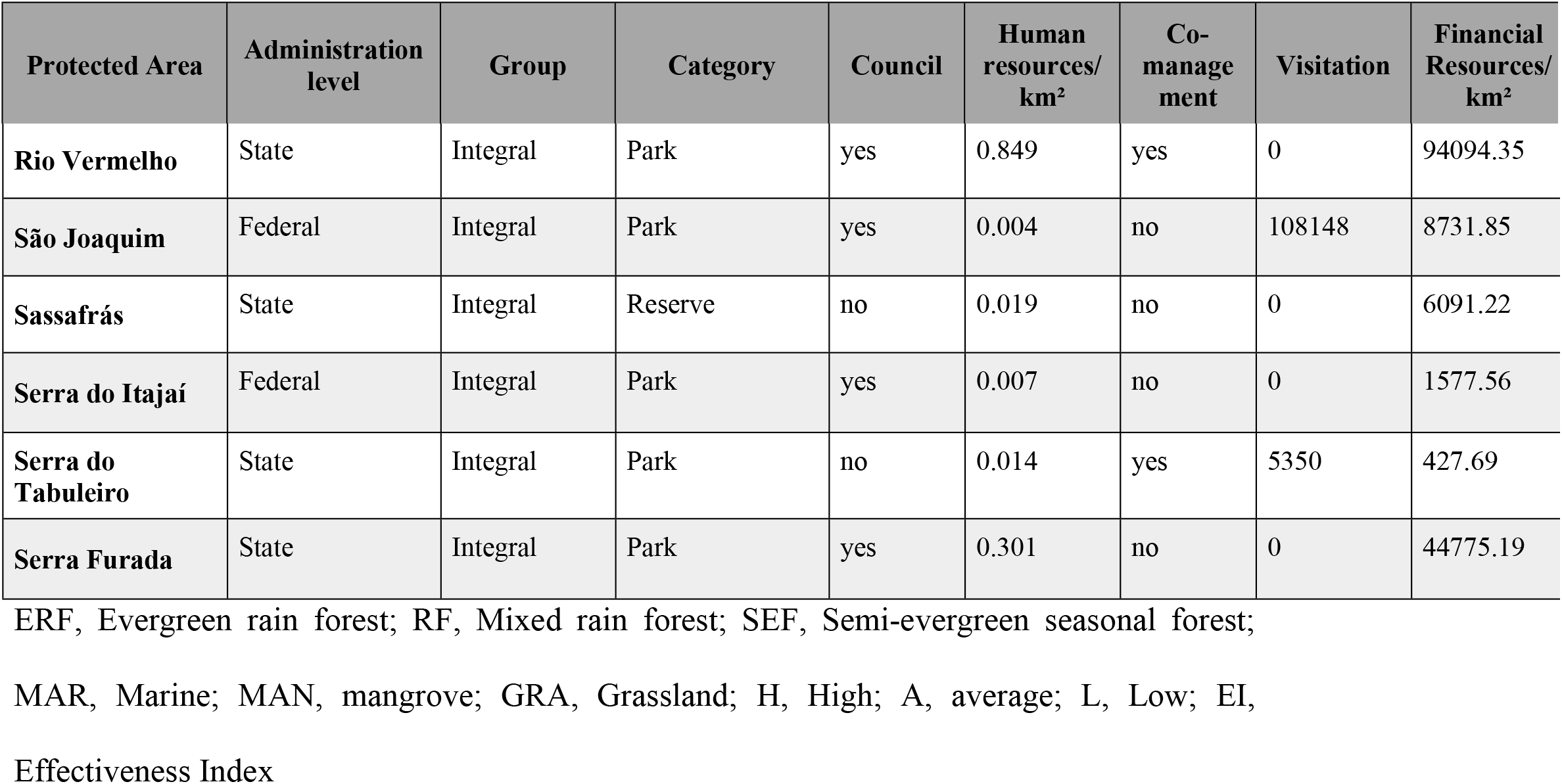
Protected areas aspects and effectiveness results.

We computed all analyses in R [24] using “vegan”, “MASS” and “car” packages.

## Results

### Overview of management effectiveness and its elements

Two thirds (14/21) of the PAs had average management effectiveness. Of the remaining, management effectiveness was high in 6/21 PAs and low in one (Fig 1). On a numerical scale, mean effectiveness index was 0.53 (min = 0.37; max = 0.75), which corroborates a medium overall management effectiveness. Under federal administration, effectiveness was about evenly distributed as either average (6/11) or high (5/11) (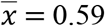; min = 0.43; max = 0.75), thus being very close to high effectiveness. On the state level, 8/10 PAs had average effectiveness, while one had high and another had low effectiveness (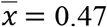; min = 0.37; max = 0.64), being closer to falling under low than under high effectiveness. Based on mean indices, federal level PAs showed an improved management effectiveness by 25% over state level PAs.

**Fig 1.**
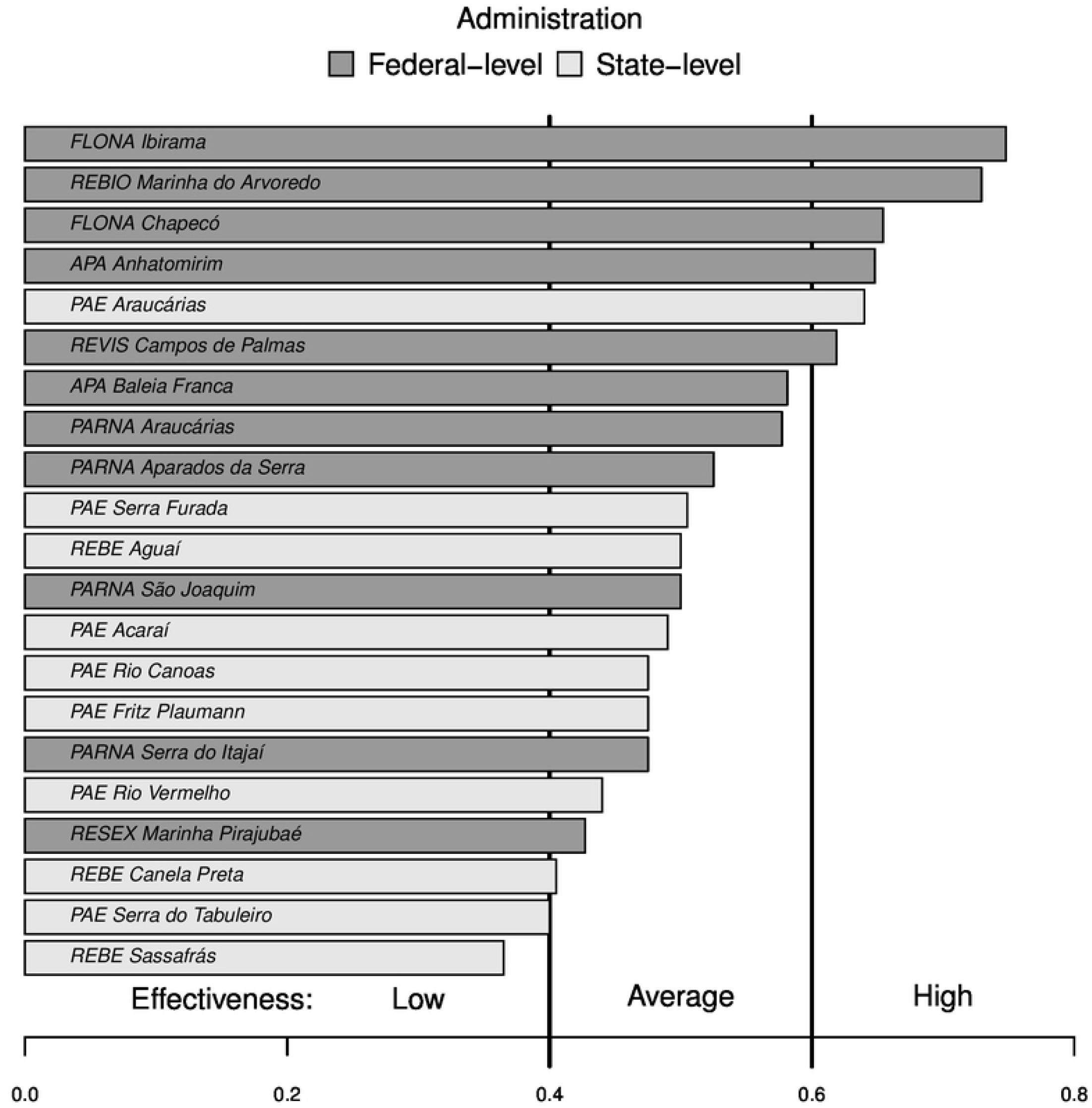
Management effectiveness of Federal-level (dark grey) and State-level (light grey) protected areas. FLONA=National Forest, REBIO or REBE=Biological Reserve, APA=Environmental Protection Area; PARNA=National Park; PAE=State Park; RESEX=Extractivist Reserve.

Context was the best managed element of effectiveness 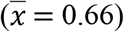, while input was lowest 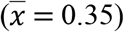 (Fig 2). A similar pattern was found for PAs from either federal level (context: 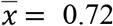; input: 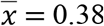) or state level administration (context: 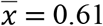; input: 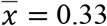). Mean scores of all elements were higher for federal than state level PAs, except for planning, which was lowest for federal level PAs.

**Fig 2.**
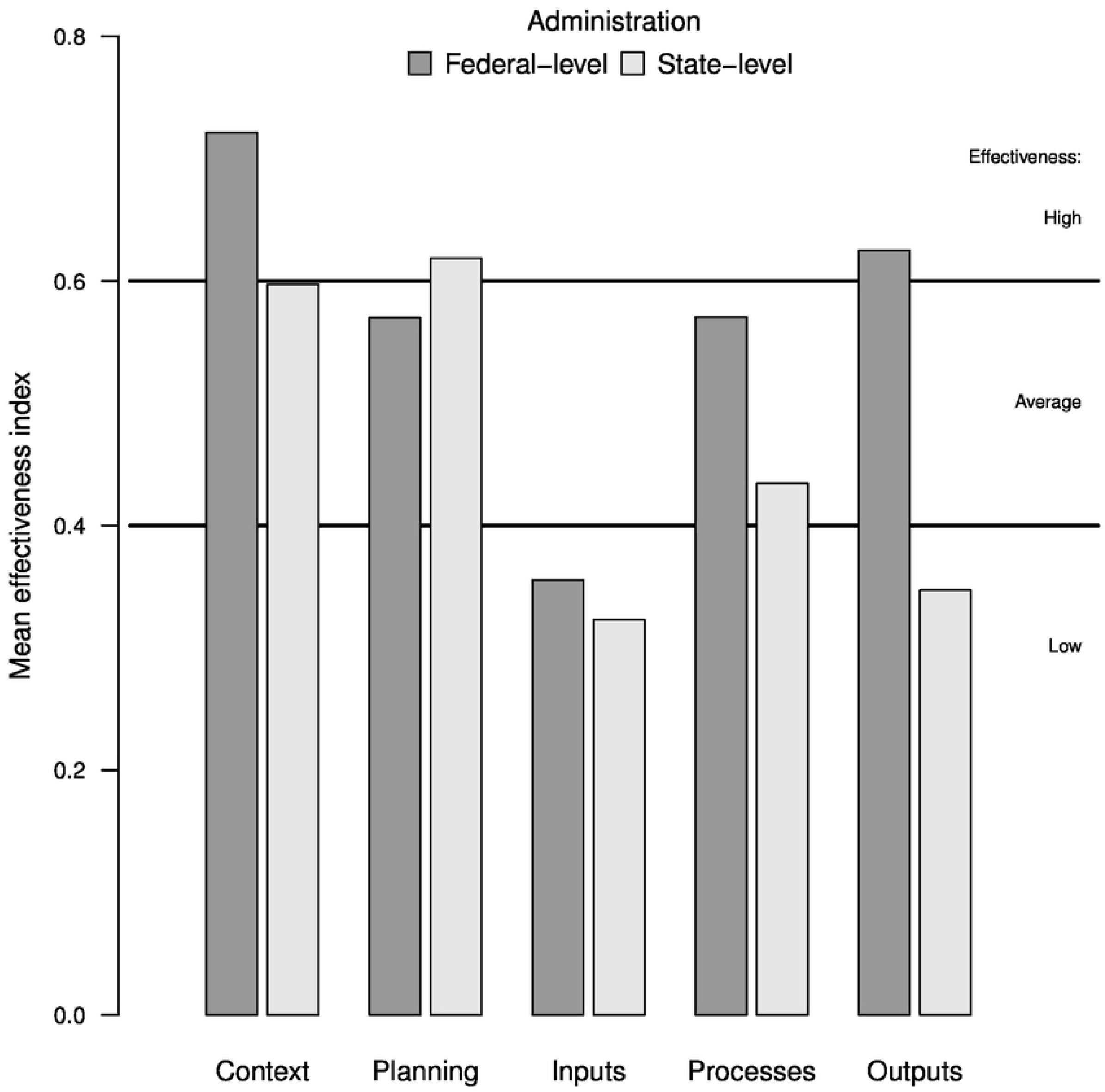
Protected areas management effectiveness per management element.

We identified 22 pressures and threats across PAs (min = 4, max = 18 per PA) that can require management actions. Criticality level ranged from 82 to 906. On average, there were ~11 pressures and threats per PA (criticality 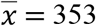). Top pressures and threats (with greater degrees of criticality) were 1) hunting; 2) invasive alien species; and 3) construction or operation of infrastructure.

### How protected areas aspects vary, and which aspects might drive successful management?

Most PAs were in the broadleaf evergreen rainforest but spanned over seven different ecosystem types (Table 1). Funds per km^2^ differed in up to 2,022 times among PAs. Only 2/21 PAs had > 1 employee/km^2^ and 13/21 PAs had < 0.1 employee/km^2^. Land tenure was resolved in 8/21 of the PAs and most PAs had management plans. Of federal level PAs, all had < 1 employee/km^2^. Regarding the protection category, 5/11 of federal level PAs were of full protection and 6/11 of sustainable use. Land tenure was resolved in 6/11 federal level PAs and 3/11 lacked management plans. Under state level administration, land tenure was resolved in 3/10 PAs and 3/10 had no management plan. All state level PAs were of full protection category.

PA aspects explained just over 50% of overall management effectiveness (adj. R^2^ = 0.52; F = 4.65; P < 0.01). The predictors kept at this stage were administration level, ongoing co-management arrangements, funds (negative relationship), management plan availability, tenure resolution, and criticality of pressures and threats (negative relationship). Because the model with the lowest AICc contained non-significant predictors, we further removed those with high P-values up to the limit of ΔAICc < 4 from the above-mentioned model. After that, administration level (P = 0.01) and management plan availability (P = 0.04) were deemed the most important aspects for management effectiveness (adj. R^2^ = 0.52; F = 4.65; P < 0.01). Adding back predictors in this model resulted in tenure resolution to be significant (P = 0.025), but with ΔAICc > 4. Thus, even though land tenure is always of concern, its effects tended to be overtaken by administration level and management plan availability.

Regarding the relationship between PA aspects and effectiveness, protection category was removed from the redundancy analysis (RDA) because of multicollinearity. After step-by-step reduction, the final RDA explained ~50% of the variation in the elements of management (Fig 3). Tenure resolution (F = 12.48, P < 0.001), administration level (F = 7.01, P < 0.01), degree of criticality of pressures and threats (F = 5.72, P < 0.01), and management plan availability (F = 3.57, P < 0.05) were the most important predictors in explaining the elements of management effectiveness, and tenure resolution was the key aspect to ensure management to be successful. PAs with tenure resolution had also the highest values for both the elements, processes, and outputs. In turn, being under federal administration was the second most important aspect for improving management effectiveness. Federal level PAs also tended to have the highest values for both context and outputs elements. Criticality of pressures and threats was the third most important aspect and had mostly an inverse relationship with effectiveness. Criticality increased towards PAs with larger values of context and lacking management plans, and away from PAs with larger values of input.

**Fig 3.**
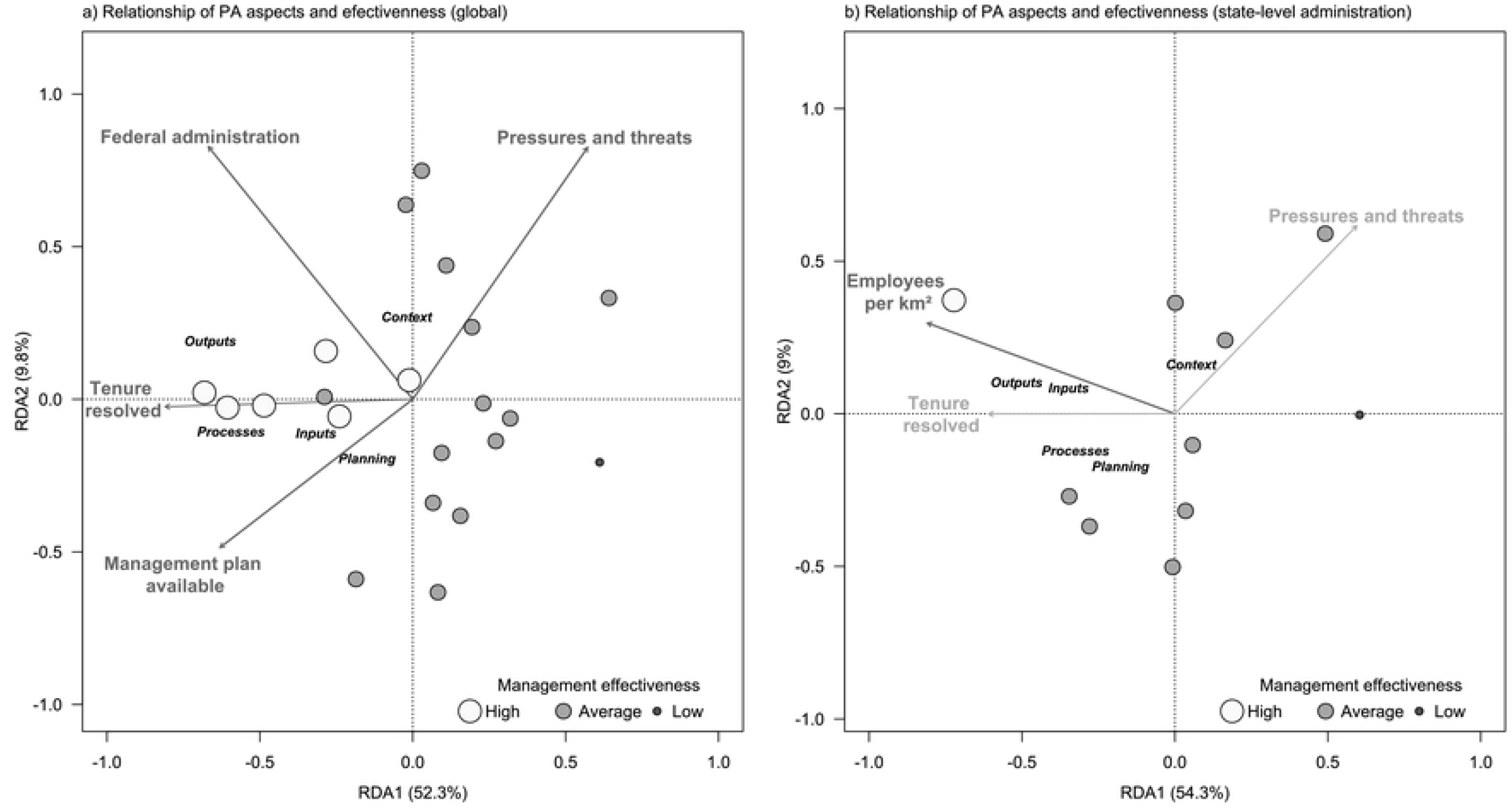
Relationship of protected areas aspects and effectiveness for (a) all areas, and (b) state-level administration.

For the state level data and after both checking for multicollinearity and model simplification, the number of employees per km^2^ was the aspect most strongly related to changes to the effectiveness index, being followed by management plan availability (Fig 3). The model with these two aspects showed a good fit to data (adjusted R^2^ = 0.74; F = 14.07; P < 0.01). When assessing changes to the elements of effectiveness, the most relevant aspects was again the number of employees per km^2^ (P < 0.01). In addition, tenure resolution and criticality of pressures and threats tended to contribute to a smaller extent. The RDA including only the number of employees per km^2^ as a predictor explained 42% of variation in effectiveness elements, whereas with the three aspects explanation reached 63%. The RDA with the three aspects was kept as to further explore the results. This RDA showed enlarging staff size leads to more successful management effectiveness in state level PAs, especially because it was positively linked to larger output value. PAs with tenure resolution had higher values for the elements of effectiveness. In turn, there was a negative relationship of elements with criticality of pressures and threats, being associated with lower values for processes and planning elements.

## Discussion

We found the protected area network to be of average management effectiveness. Some aspects of PAs postulated to affect management effectiveness were found relevant. Our main hypothesis was rejected because staff and fund sizes were poor drivers of successful management. Instead, land tenure issues seem to be the top barrier for more successful management. Administration level, development of management plans, and pressures and threats were additional aspects associated with differences in management effectiveness.

The average effectiveness corroborates global [10] and federal level PAs assessments in Brazil [23]. Context of PAs was the element of management with the highest values, indicating improvement in prioritization schemes and planning before creating PAs [25]. Elements processes and outputs showed a high relationship with effectiveness. We attribute this result to the practical characteristic of these elements, because processes comprise the execution of the management actions, while outputs gauge action results [8]. The element with the lowest values was inputs, pointing to a shortage of resources to carry out management actions [9].

PAs under federal administration were more effective than state level PAs. Such results can be due to 1) recurrent effectiveness assessments [23]; 2) structure of the management body; and 3) programs developed at the federal level. Federal level PAs have already gone through three different assessments and effectiveness indices improved along them, pointing to benefits along the process. The same pattern showed up worldwide, where ~70% of PAs increased effectiveness in reassessments [26]. In Brazil, the federal management body used to be more structured [27], with programs developed at this level often maintained with international resources. Still, results for PAs of the State of Santa Catarina are above the average of other Brazilian states (e.g. [28]), suggesting barriers to be distinct or more detrimental in other PA networks.

Resolving land tenure issues has already been identified as a priority [19], but see also [10]. We reinforce tenure resolution is crucial for three reasons. First because the creation of PAs without compensation establishes a conflict between the right to property and the right to a balanced environment in the Brazilian constitution. Second, activities incompatible with conservation aims can proceed until tenure is granted, pressing the PA and making management actions more difficult [29]. Third, tenure issues can trigger other conflicts [30] because of the large involved figures, administrative difficulties, and fragility of land tenure structure [31]. However, these barriers should neither inhibit creation of new PAs, be an excuse for the use of less restrictive categories, nor restrict new PAs to public land to avoid acquisition [32].

The relationship found between pressures and threats with context is worrisome. Indeed, PAs under highest conservation priority given their biodiversity tend to be under more pressures and threats. Although hunting and deforestation are top threats to PAs globally [5, 17], deforestation was irrelevant here. Hunting, conversely, was identified as critical and is of concern because it has direct (population reduction or extinctions, [33]) and indirect effects on biodiversity (changes on either community structure and ecosystem functioning, [34]; animal behavior, [35]; or plant dispersal and regeneration, [36]). Combating hunting is challenging because of its stealth aspect and because counteracting actions often need to go beyond PA limits. Another critical threat we found were invasive alien species, which reduce diversity and can alter the structure and the functioning of ecosystems (e.g. [37]). Costs of controlling invasive species are among the main management obstacles for PAs [38]. Infrastructure is also among the most significant pressures with direct impacts on PAs. Roads lead to roadkills [39], ease deforestation [40] and dispersal of invasive alien species. Reducing infrastructure impacts requires integration of environmental policies and improvement of environmental impact assessments and mitigation [41]. Such PAs require both more resources, especially employees [18], and meaningful management plans. However, the current situation is worsening, following pressure to simplify the assessments [42], evidence of corruption [43] and strong and coordinated political interference in the decision-making process [44].

Management plan development was one of the main aspects associated with differences in effectiveness, pointing out they are important and likely also meaningful for management [15]. Management plans precede execution of actions and define guidelines for optimization of resources. Although such precedence is both obvious and plan development considered urgent under current legislation, many PAs still lack such plans, thus contributing to poor management effectiveness. Moreover, we found little relevance of staff and fund sizes. Although both are often bottlenecks for conservation [14] and found here to match low values of the input element, all these aspects were unrelated with effectiveness. We also expected co-management to bring benefits by improving staff size [13] and visitation to increase with management effectiveness [16], but such relationships were rejected, indicating that changing how co-management or visitation occurs is necessary to result in gains for PAs [45]. Likewise, having an instituted board seems insufficient to ensure proper representation and governance and, thus, be ineffective on management effectiveness (but see e.g. [12]).

## Conclusions

The protected area network we assessed has currently an average effectiveness. This means they are just partially fulfilling their objectives. We identified four main aspects to explain the differences in effectiveness: management plan availability, current and potential impacts caused by threats, tenure resolution; and administration level. Except for being under federal level administration, these aspects are related to actions and measures that can be addressed by management actions and by a greater alignment with the legislation related to protected areas, which demonstrates its low application. We suggest the systematization of these aspects in a management model based on three top priorities: 1) development of management plans because of its importance in establishing guidelines for the other aspects; 2) management actions to eliminate or reduce the effects of the most important threats (hunting, invasive alien species, and installation and maintenance of infrastructures); and 3) land tenure resolution, which seems even more strongly related to effectiveness than the priorities above. As a cautionary, tools such as RAPPAM are likely to fail if used as the sole gauge of PA management effectiveness and any benefits of assessments can be easily lost following political neglect. Minding such limitations, we suggest a great opportunity to improve management effectiveness of PAs by means of recurrent assessments. Only in this way will it become clear when a barrier to an effective management has been cleared and which one is to be tackled next.

## Acknowledgements

We thank all people who provided information about state-level PAs and to ICMBio for allowing access to RAPPAM data for federal-level PAs. Special thanks to the managers of protected areas for their dedication to this challenging work. MEM thanks to IMA (Instituto do Meio Ambiente de Santa Catarina) for a partial release to his Master’s course

